# Topographic effects on dispersal patterns of *Phytophthora cinnamomi* at a stand scale in a Spanish heathland

**DOI:** 10.1101/281691

**Authors:** Enrique Cardillo, Angel Acedo, Enrique Abad

**Affiliations:** Instituto del Corcho, la Madera y el Carbón Vegetal, Centro de Investigaciones Científicas y Tecnológicas de Extremadura (CICYTEX), Mérida, Spain; Departamento de Física Aplicada and Instituto de Computación Científica Avanzada (ICCAEX), Universidad de Extremadura, Mérida, Spain

**Keywords:** epidemiology, *Erica umbellata*, disease spatial pattern, topography

## Abstract

*Phytophthora cinnamomi* is one of the most important plant pathogens in the world, causing root rot in more than a thousand plant species. This observational study was carried out on a *P. cinnamomi* infected heathland of *Erica umbellata* used as goat pasture. The patterns and shapes of disease foci and their distribution were described in a spatial and temporal context using an aerial photograph record. A set of topographic traits was selected on the basis of a disease dynamic hypothesis and their effects on observed spatial disease patterns were analyzed. Incipient infections situated in flat terrain expanded as compact circular front patterns with a low growth rate. On slopes, disease patches developed more rapidly down slope, forming parabolic shapes. The axis direction of the parabolas was highly correlated with terrain aspect, while the parabolic amplitude was associated with land curvature and slope. New secondary foci appeared over the years producing an accelerated increase of the affected surface. These new foci were observed in sites where disease density was higher or near sites more frequently visited by animals such as the stable or the forage crop. In contrast, a smaller number of disease foci occur in areas which animals are reluctant to visit, such as where they have a short range of vision. Our results suggest that 1) the growth of existing *P. cinnamomi* foci is controlled by a combination of root-to-root contact and water flows, 2) the increase in the diseased area arises mainly from the multiplication of patches, 3) the formation of new foci is mediated by long-distance transport due to the movement of animals and humans along certain preferential pathways, and 4) geomorphology and topography traits are associated with the epidemiology of this soil-borne pathogen.

## Introduction

Many forest and agricultural ecosystems exposed to the impacts of one of the world’s most invasive species, *Phytophthora cinnamomi* [1]. *Phytophthora cinnamomi* is a widespread soil-borne pathogen that infects more than one thousand woody species, causing root rot and especially impacting countries with Mediterranean climate [2]. This plant disease has been reported as the causal agent of the *Phytophthora* dieback epidemic in the jarrah (*Eucalyptus marginata*) forest of south-western Australia [3], and of the massive decline of cork and holm oak in south-western Europe [4, 5]. It is also involved in diseases affecting fynbos vegetation in the Cape region of South Africa [6] and oak in California [7]. *Phytophthora cinnamomi* causes major losses to other forest systems such as chestnuts [8, 9] and horticultural trees such as avocados [10].

To gain knowledge for disease management, spatial analysis of dispersal patterns can provide information on inoculum sources, transport, release mechanisms and disease-conducive environments. Ultimately, spatial patterns of plant disease invasion are the consequence of the effects of environmental forces on dispersal processes [11]. Therefore, studies on pathogen or disease distribution contribute to identifying key epidemiological factors and predicting future disease scenarios by modelling. However, the insights obtained are strongly dependent on the specific spatial scale, since the choice of scale may limit our ability to understand certain aspects of the epidemic, but also facilitate the interpretation of other features [12]. Prior to the present work, global- and regional-scale distribution models have been employed to map current and future potential ranges of *P. cinnamomi*. Models at these scales are invaluable to assess not only the impact of future climate scenarios [2] on the risk occurrence of this disease, but also the combined effects of major factors in this context, such as soil humidity and temperature [13, 14]. Additionally, analysis at these scales has revealed the importance of *Phytophthora* dispersal paths such as international trade in plant material and tourism [15].

At a landscape scale, other risk factors such as forest fragmentation, habitat heterogeneity or geophysical features could emerge [16, 17]. For example, a shorter distance to the forest edge resulted in an increased mortality risk associated with the sudden oak death disease that *Phytophthora ramorum* caused in California [18]. Working at a landscape scale, Keith et al. [19] recognized the effect of distribution and abundance of susceptible host plants on *P. cinnamomi* location patterns. In Iberia, Moreira and Martins [20] and Costa et al. [21] highlighted the role of the presence of shrubs in oak decline. At this level of spatial hierarchy, human and vehicular traffic [22] as well as the movement of domestic and wild animals [22, 23, 24; 25] have been identified as propagule vectors.

At a stand scale, which is usually the most relevant to disease managers, *P. cinnamomi* is able to use several mechanisms of dispersal: soil spread, root to root contact, water transport and dispersal by humans and animals [11, 26, 28]. Nevertheless, the observation and understanding of dispersal mechanisms at play in a given site is not easy because the precise location of inoculum sources, transport paths and sinks is hard to determine. In the Bassendean Dune ecosystems of Western Australia, observation of infected Banksia woodlands by aerial photographs revealed a slow rate (about 1 m/year) pattern of disease spread. Several authors [26, 27, 28] have described the shapes of these patterns and hypothesized that root-to-root contact was the predominant mechanism of invasion.

Although it was known that *P. cinnamomi* propagules can be dragged by water more than 120 m downslope through some soils [29], and that the propagules may travel even further when being carried by water flows [22, 3], our understanding of soil water transport is far from complete. Some studies report a general association of pathogen presence with water abundance in the soil. For example, in a specific area of southern Ohio, *P. cinnamomi* was more frequently isolated in moist sites of white oak (*Quercus alba*) forest [30]. In another study on the spatial distribution of chestnut ink disease (*P. cambivora*), a negative relationship was found between disease incidence, severity and tree mortality on the one hand and distance from drainage on the other [9].

In the above context, the propagation distance measured from inoculum sources is considered to be a general and important factor of infection risk. In this sense, clustered patterns of the pathogen have been frequently observed in the vicinity of symptomatic trees [31], but also near asymptomatic vegetation [30, 32]. A current research topic concerns the determination of the shape and density distribution of such clusters. In this context, a highly interesting question is whether such features can be successfully reproduced and predicted by dispersal models relying on reaction-transport equations [33]. In this respect, topographic features influencing the local distributions are likely to play an important role.

Historical collections of aerial photography and modern high resolution imagery have proven to be an invaluable resource to detect, quantify and map *Phytophthora* diseases of forest trees [26, 34, 35]. Nevertheless, relevant environmental data such as soil moisture, temperature and fertility or information about human, animal or water flows are more difficult to obtain at a resolution matching the granularity of a pathosystem at stand scale. To fill this gap, primary geo-morphologic attributes such as altitude, slope and aspect, or derived attributes such as topographic curvature and radiation indices have been extracted from high resolution digital terrain models [36, 37]. These variables have been used as proxies of environmental factors underlying the biological processes and the spatial distribution of *Phytophthora* pathogens [9, 19, 20, 27]. The same approach can be used to estimate the concentration or density of animals acting as propagation vectors. It has for instance been shown that, besides forage quantity and quality, the distribution of large herbivores is affected by abiotic factors such as water availability, weather, and topography [38, 39]. In this sense, key topographic traits such as aspect, slope, solar radiation or visibility range have been reported to be habitat selection features [39, 40, 41].

In more general terms, the epidemiological analysis of infectious diseases in plant areas aims to describe, understand, compare, and predict epidemics [42]. Mathematical and statistical models, be they static or dynamic, are used to reduce the epidemic phenomenon to its essential characteristics and to describe the main mechanisms underlying the progress of the disease. More specifically, dynamic approaches aim to reproduce the time evolution of the epidemics by means of a set of differential equations that are in general coupled to one another. Such differential equations contain deterministic and/or stochastic terms mimicking the effect of the most relevant factors behind the spread of the infection. Beyond conventional approaches such as the healthy-latent-infectious-removed model [42], the spread of plant diseases and biological invasions in general are increasingly being analyzed and simulated with the help of models largely inspired by the theory of reaction-diffusion systems.

In their simplest form, reaction-diffusion models pair two ingredients, namely a population growth function and the dispersal kernel describing transport via Fickian diffusion. In this simple setting, the invasion progresses along concentric, normally distributed, frontal waves whose propagation velocity depends on the diffusivity of the system [33]. Additional terms can be incorporated to the relevant equations in order to reflect more complex situations such as directional dispersion (advection). A representative example is given by a field of wind force influencing the dispersive transport of *Phytophthora infestans* spores [33]. In other cases, the diffusion kernel could be adapted to describe a stratified or dual mechanism of dispersion consisting of concurrent processes of neighbourhood and long-distance spread [43] that conspire to accelerate epidemic spread [44]. In any case, a realistic description should account for the coupling between the dynamics of the pathogen and those of the ecological drivers of the infection, whereby the stochastic nature of certain variables cannot be safely ignored in general. Thompson et al. [14] adopted this approach to address multiple pathways by which climate (soil humidity and temperature) can constrain the dispersal range of *Phytophthora cinnamomi* at a regional scale.

Such models are ideal cause-effect simulators; however, identification, understanding and simplification of the real processes involved are required in order to obtain a tractable system that can be solved analytically or numerically. Therefore, a first step prior to the development of more sophisticated models is the use of non-mechanistic models to gain some preliminary insights and to develop an intuition for the behaviour of the system at hand. In this case, several snapshots reflecting the development of the infectious disease are used to extract spatial data of the progress of the disease that are subsequently related to environmental variables by parametric or non-parametric methods [34]. Some non-parametric models, such as maximum-entropy methods, allow the modelling of biological distributions from the inferred presence of the pathogen alone [45]. However, when both types of datasets are available, namely spatial distributions for the pathogen and the environmental variables, case-control studies are widely used in epidemiology to retrospectively analyse the effect of explanatory features on disease events. To quantify the relationship between risk factors and disease status, odds ratios (relative risk) and their standard errors are estimated, typically via logistic regression models. Case-control designs and logistic regression have been used in *Phytophthora* forest disease studies to demonstrate the relationship between the crown status of trees and the presence/absence of the pathogen [46], to identify environmental risk factors [27], or to investigate whether the characteristics of the trees and/or stand facilitate the progress of the disease [47].

Our team has recently reported how *P. cinnamomi* causes massive mortality among populations of dwarf heath (*Erica umbellata*) in southwestern Spain [48]. In this study, roots and rhizosphere soil samples from *E. umbellata* plants exhibiting dieback and mortality were collected from 11 foci. A *Phytophthora* sp. was consistently isolated from 73% of soil samples and from 46% of roots. All isolates were identified as *P. cinnamomi* by morphological and cultural traits and ITS region sequencing. During the pathogenicity test conducted to verify the causal agent, all plants inoculated with *P. cinnamomi* died within just 14 days of pathogen infection, revealing a very high susceptibility to the pathogen. Moreover, the spread of the disease in a monospecific heathland was easily observable, since it formed solid foci delimited by a neat and narrow front of dead and dying heaths. Therefore, an excellent opportunity emerged to describe the spread of *P. cinnamomi* at high spatial resolution, to analyse the associated factors, and to provide a first approximation to a theoretical description in terms of simple fitting models.

This work aims to gain understanding of the spread patterns of *P. cinnamomi* at stand scale by combining aerial imagery, topography, geomorphology and field data of a heathland in Spain. Our main focus will be on four tasks, namely: 1) establishing a causal relationship between the observed disease fronts and the presence of *P. cinnamomi,* 2) describing the observed patterns quantitatively and making hypotheses on probable dispersal mechanisms, 3) analyzing the relationship of the latter with terrain morphology and topography, and 4) gaining some insight into the relative importance of the different dispersal paths.

## Materials and methods

### Area of study

The study was carried out on *Erica umbellata* heathland located on the quarzitic mountain range ‘Sierra de las Villuercas’, in eastern Extremadura. This mountainous area (1,500 km^2^), is part of Montes de Toledo, a large east-west mountain system crossing the Spanish central plateau. Villuercas has a Mediterranean climate with dry summers and mild and wet winters. Mean annual rainfall is around 880 mm, but only 3 mm in July. Mean temperature in July is 27.2 ^º^C and 6.4 ^º^C in January (averages in the period: 1980-2010). Soils are glacis-piedmont surfaces dating from the middle to upper Pliocene and locally known as ‘rañas’ [49]. They are very acidic (pH 5.0) and poor quartzitic Ultisols with bad drainage.

The 200 ha study site (Fig. 2), known as ‘Las Alberquillas’, is located 10 km south of Guadalupe, in the province of Cáceres. Its geographic coordinates are 39^º^ 21’ 46’’ N and 5^º^ 19’ 36’’ W (ETRS89) and the altitude is 650 m. The general slope is oriented to the southwest with a gentle gradient (5%). However, this easily eroded terrain is crossed by ephemeral streams where the slope gradient increases abruptly (> 50%). The study area is populated by a monospecific (95% cover) shrubland of dwarf heather (*Erica umbellata*). Other shrubs, such as *Halimium ocymoides* and *Pterospartum tridentatum*, are sparsely present. The surrounding landscape is composed of natural cork-oak forests (*Quercus suber*), and forestry stands comprising maritime pine (*Pinus pinaster*), and eucalyptus (*E. camaldulensis*).

Marginal cereal (rye) cultivation was practiced in the area after the Spanish Civil War (1940-50) [50]. The heathland is now mainly used for goat pasture, with small areas seeded with rye as fodder. In the Villuercas area, there were 57,619 goats in 2007 browsing in flocks of 260 individuals per holding on average [51]. Wild boar, deer, cattle, and sheep are present in the heathland sporadically. Patches of dead heath (Fig. 1) were first observed in the area in spring 2011.

**Fig. 1.**
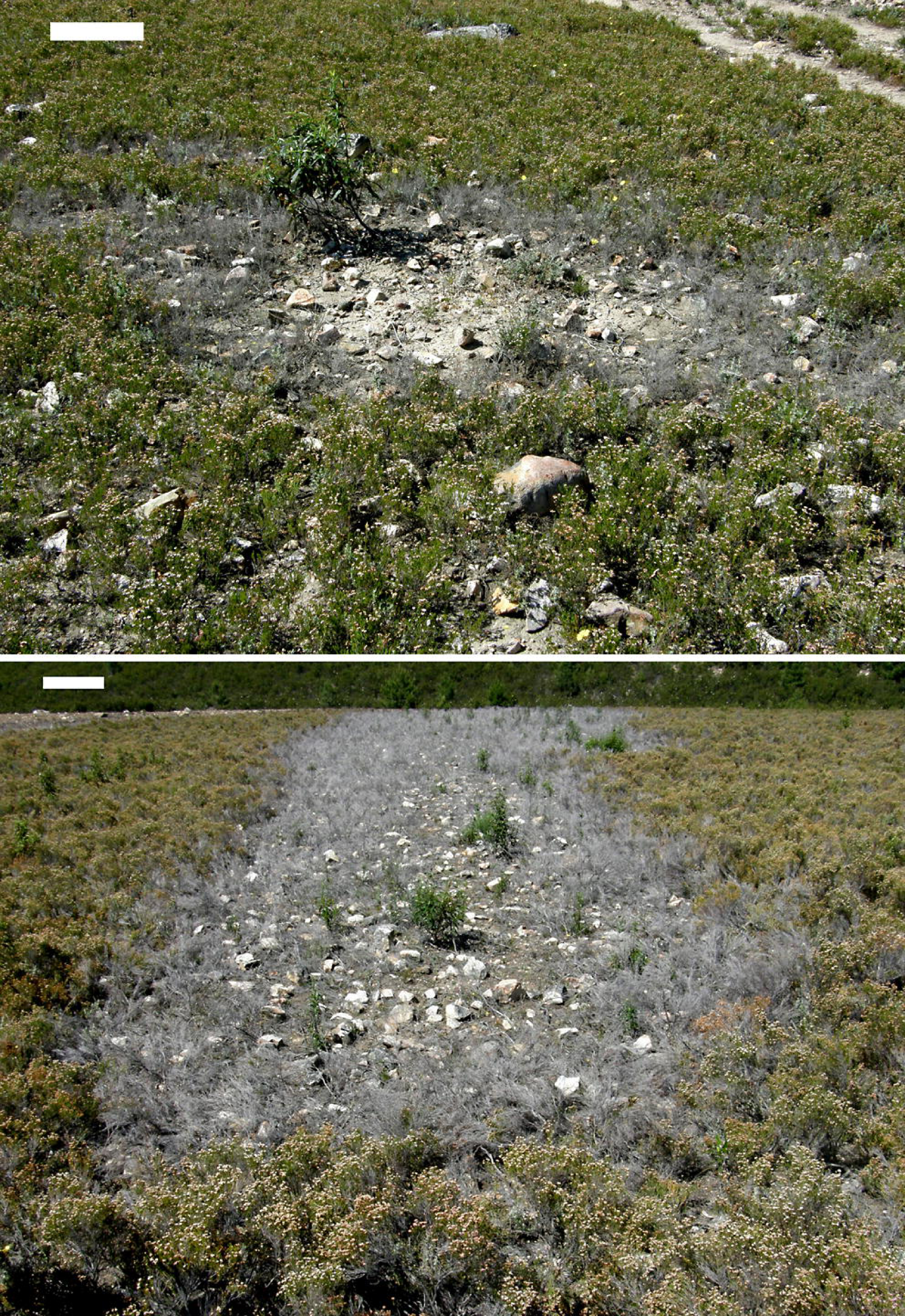
**Patches at initial stage of *Phytophthora cinnamomi* infection on a heathland of *Erica umbellata*. Top:** small focus with a circular shape on a flat area. **Bottom**: a focus spreading downhill on a steeper slope. Colonization of the bare soil by *Cistus ladanifer* started inside the disease front. White bar at the top left is approx. 50 cm length. Images taken by the authors.

**Fig. 2.**
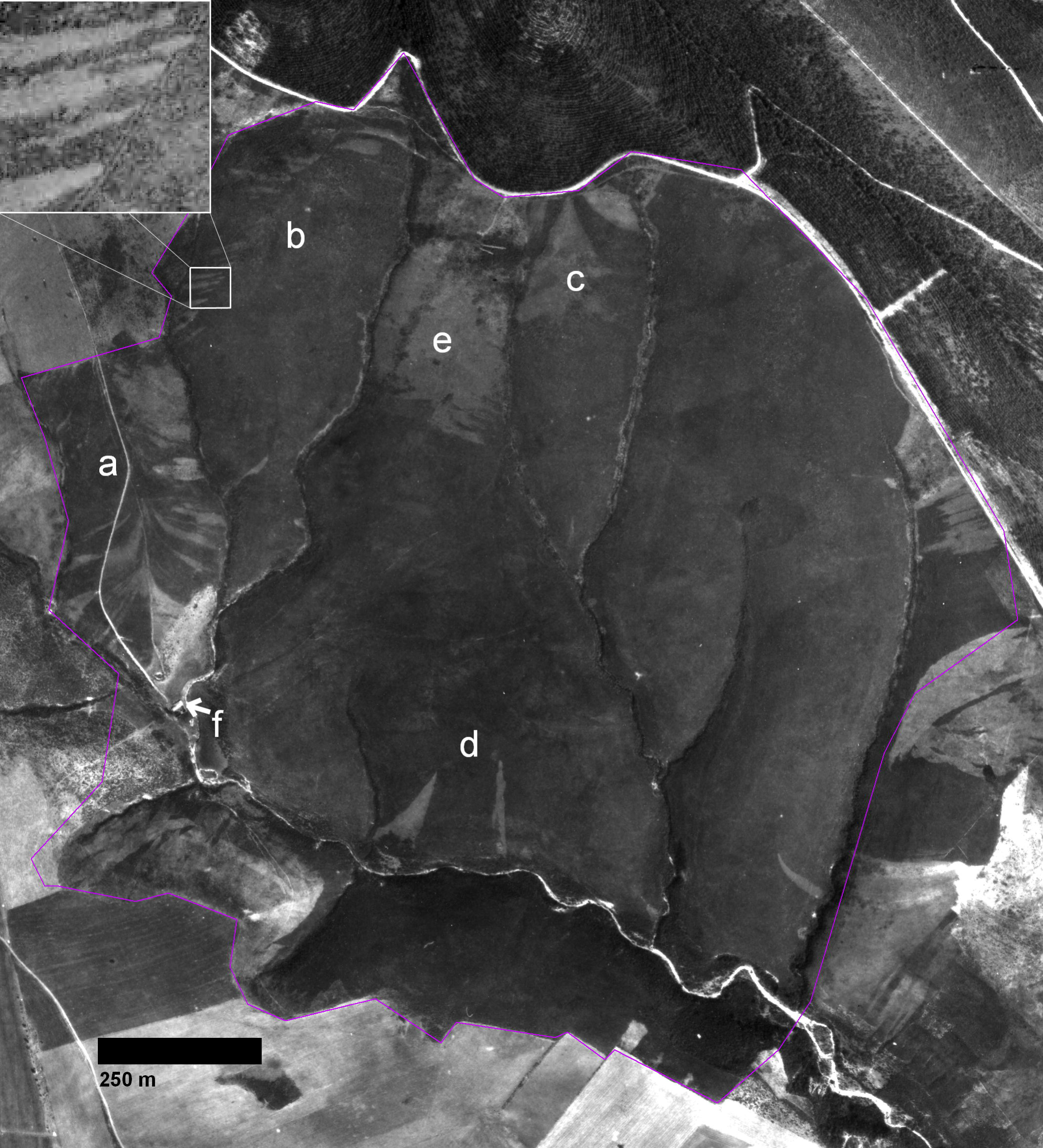
**Panchromatic aerial image of the study area photographed in 1981**. Triangular patches of pale grey colour are disease foci originated by *Phytophthora cinnamomi* infection of *Erica Umbellata* heathland (dark grey background). Foci can be seen emerging from the farm access road (**a**), tractor tire tracks, see the upper left magnification (5x) window (**b**), main road at the north edge (**c**), or a few dispersed points in the south (**d**). A cultivated rye paddock (**e**), and a goat stable, marked with a white arrow (**f**) are identifiable too. Purple line shows study area limits. Printed under a CC BY license, with permission from Junta de Extremadura CICTEX, original copyright: Orthoimagery CC-BY 4.0 CICTEX, Junta de Extremadura.

### Pathogen isolation

To confirm that *P. cinnamomi* was not extensively present outside diseased patches and the symptomatic heath was a reliable proxy for the spread of the pathogen, a survey was conducted. In this survey, a total of 18 samples were collected along three horizontal transects radiating outward from the patch centres. Six soil samples were collected per transect: two (50 cm apart) in the bare ground of the patch core, two (20 cm apart) in the disease front with dead and symptomatic heaths and two more just outside (50 cm apart) the patch among healthy vegetation. Each sample, 100 ml of soil extracted from the rhizosphere, was baited using ten young cork oak leaves [52]. After 48 hours, each necrotic spot on the leaves was plated onto NARPH agar [66]. The soil pathogen load was assayed as the number of *P. cinnamomi* positive isolations on leaves. A general linear model (GLM), using Poisson response and log link, was applied to the obtained set of numbers. This was done by considering the effect of the location of each sample site, categorised as being inside, at or outside the disease front. The mean number of isolations in every position was compared using Tukey’s HSD test.

The field study was carried out on private land under permission of its owner. No endangered or protected species were involved and all plant samples were gathered under supervision of a forest ranger from the environment agency of local government (Junta de Extremadura).

### Imagery (1956-2012) and geomorphologic data

Over the study area, a time series (1956-2012) of aerial photographs was gathered from the Cartographic and Territorial Information Centre of Extremadura. The set consisted of the imagery (Table 1) from the Spanish National Program of Aerial Orthophotography and historical images taken by other agencies (Orthoimagery CC-BY 4.0 CICTEX, Junta de Extremadura). Some of them were orthorectified in order to achieve geographical matching and measurement accuracy. All images were enhanced to improve visual quality.

**Table 1.** **Aerial imagery collections used in this study**

| *Date* | *Agency* | *Technology* | *Spectral resolution* | *Scale* | *Spatial resolution* |
| --- | --- | --- | --- | --- | --- |
| 1956 June | US Army Map Service | Film | Panchromatic, B&W | 1:33,000 | 1.00 m |
| 1981 March | MOPU | Film | Panchromatic, B&W | 1:18,000 | 0.50 m |
| 1984 August | IGN | Film | Panchromatic, B&W | 1:30,000 | 1.00 m |
| 1997 | MAPA | Film | Panchromatic, B&W | 1:40,000 | 1.50 m |
| 2000 | IGN | Film | Colour, RGB | 1:40,000 | 1.50 m |
| 2006 | PNOA | Digital metric camera | Colour, RGB | 1:30,000 | 0.50 m |
| 2009 | PNOA | Digital metric camera | Colour, RGB | 1:30,000 | 0.50 m |
| 2010 | PNOA | Digital metric camera | Colour, RGB+IR | 1:15,000 | 0.25 m |
| 2012 July | PNOA | Digital metric camera | Colour, RGB | 1:30,000 | 0.50 m |

A digital elevation model (DEM) of five-meter pixel size, generated by automatic correlation and interactive stereoscopic debugging of high-resolution PNOA images, was provided by the Instituto Geográfico Nacional (IGN, Spain). Topographic and geomorphologic attributes of primary and secondary order (Table 2) were extracted from the images or derived from the DEM using implemented algorithms of ArcGIS 10.1 extension Spatial Analyst [53] and SAGA-GIS modules [54].

**Table 2.** **Topographical attributes used as predictive variables in this study**

| Attribute | Description | Units | Range |
| --- | --- | --- | --- |
| Inoculum density | Density of foci observed in 1981 | Foci / km<sup>2</sup> | 1.6 – 38.6 |
| Distance to 1981 foci | Distance to disease foci found in 1981 imagery | Meters | 0 – 395 |
| Distance to drainage | Distance to drainage streams ephemeral or not | Meters | 0 – 226 |
| Distance to goats stable | Distance to goat livestock building | Meters | 86 – 1180 |
| Distance to forage crop | Distance to rye crop to feed the goats by browsing | Meters | 5 – 983 |
| Distance to roads | Distance to the next road | Meters | 0 – 723 |
| Terrain elevation | Height above reference ellipsoid (ETRS89) | Meters | 587 – 674 |
| Slope | Rate of maximum change in height from each cell | Percent | 1.4 – 53.7 |
| Aspect | Direction of slope from each cell | Degrees | 0 – 360 |
| Plan curvature index | Contour curvature at each cell center | Dimensionless | -1.15 – 0.71 |
| Positive openness index | Visibility from each cell (see Yokoyama et al. [55]) | Dimensionless | 1.15 – 1.57 |
| Spring solar radiation (potential) | Potential solar radiation income from March 1 to May 31 | kWH/m<sup>2</sup> | 346 – 458 |
| Topographic wetness index | Accumulated number of cells upstream of a cell | Dimensionless | 3.9 – 8.1 |
| Vertical distance to water streams | Height above the next water stream, ephemeral or not | Meters | 0 – 42 |

### Disease foci recognition

Disease fronts were detected in aerial images by tone and texture differences between dead and healthy areas. Discrimination between *Phytophthora* infected patches and other perturbations (ploughed areas, fire-breaks, tracks) was done by field visits to confirm the presence of a band of decaying vegetation along the edges (Fig 1). A disease patch or focus (used as synonymous hereafter) of *P. cinnamomi* was defined according to Shearer et al. [28] as the “area of disease expression around the point of initial inoculum introduction, with the outer perimeter usually delimited by a disease front of dead and dying infected vegetation”. Taking advantage of the presence of patches at different stages of development, the vegetation dynamics following *P. cinnamomi* invasion were inferred.

### Focus growth patterns

In order to estimate the rate of radial growth of individual foci, measurements were taken in small and more developed foci. Small patches were measured in the field and in orthophotos taken in 2010 and 2012. At initial stages they were too small to be identified and measured using low spatial and spectral resolution images taken before 2010. An annual average growth rate was estimated in three directions (upslope, downslope and contour) using measurements of 22 of these incipient patches. The shape of the foci was described using aerial imagery and photos taken on the ground.

Another sample of 15 well-developed disease patches was selected randomly among those observed in the most recent (2005-2012) and accurate orthophotos. Disease fronts were vectorized with the help of ArcGIS editing tools. Since patch shapes appeared visually lengthened downslope (Fig. 3), an inner line representing the main direction of focus spread was determined as follows. At every five meters from the upper vertex of the focus, the points situated in the middle of the horizontal distances between focus fronts were identified and joined by segments (Fig. 4). Given that, in every case, these main axes appeared to follow the slope aspect (i.e. the steepest slope path), a linear regression model, fit by maximum likelihood methods, was used to test this hypothesis (R’ statistical software version 3.1.1, [56]). The azimuth of line segments was considered to be the independent variable, and the aspect of terrain at the segment start point as the predictive factor.

**Fig. 3.**
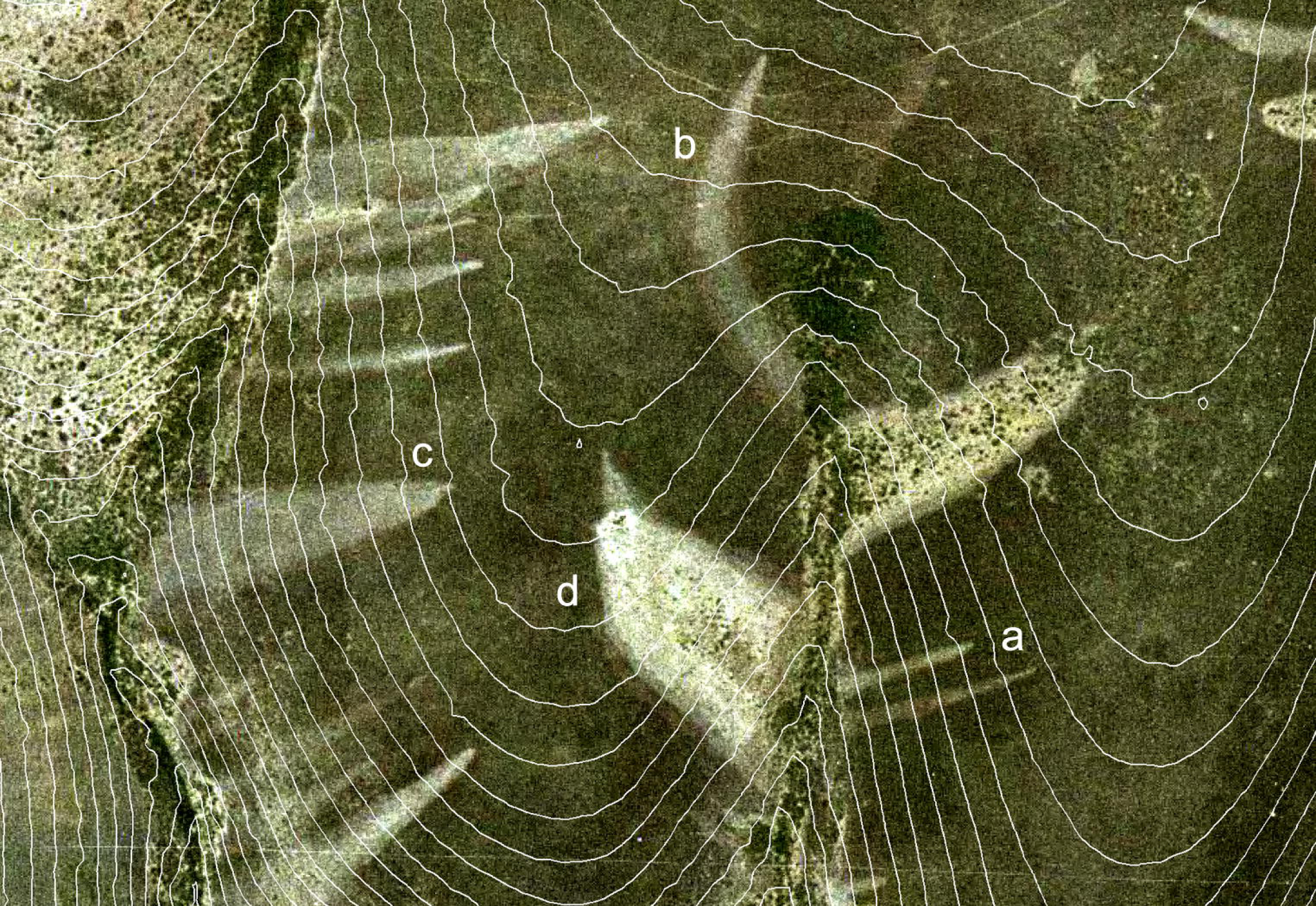
**Colour aerial image taken in 2000 showing *Phytophthora cinnamomi* infection patches on *Erica umbellata* heathland in Villuercas Sierra (Spain)**. Following traits are identifiable: needle shapes of foci at their initial stage (**a**), bending effect due to changes in terrain aspect (**b**), a wide patch in a zone with divergent curvature (**c**), well developed focus with asymmetric branches following slope aspect (**d**). All of the disease fronts have parabolic shapes ending where drainage is reached. Contour lines were superimposed to show terrain morphology. Printed under a CC BY license, with permission from Junta de Extremadura CICTEX, original copyright: Orthoimagery CC-BY 4.0 CICTEX, Junta de Extremadura.

**Fig. 4.**
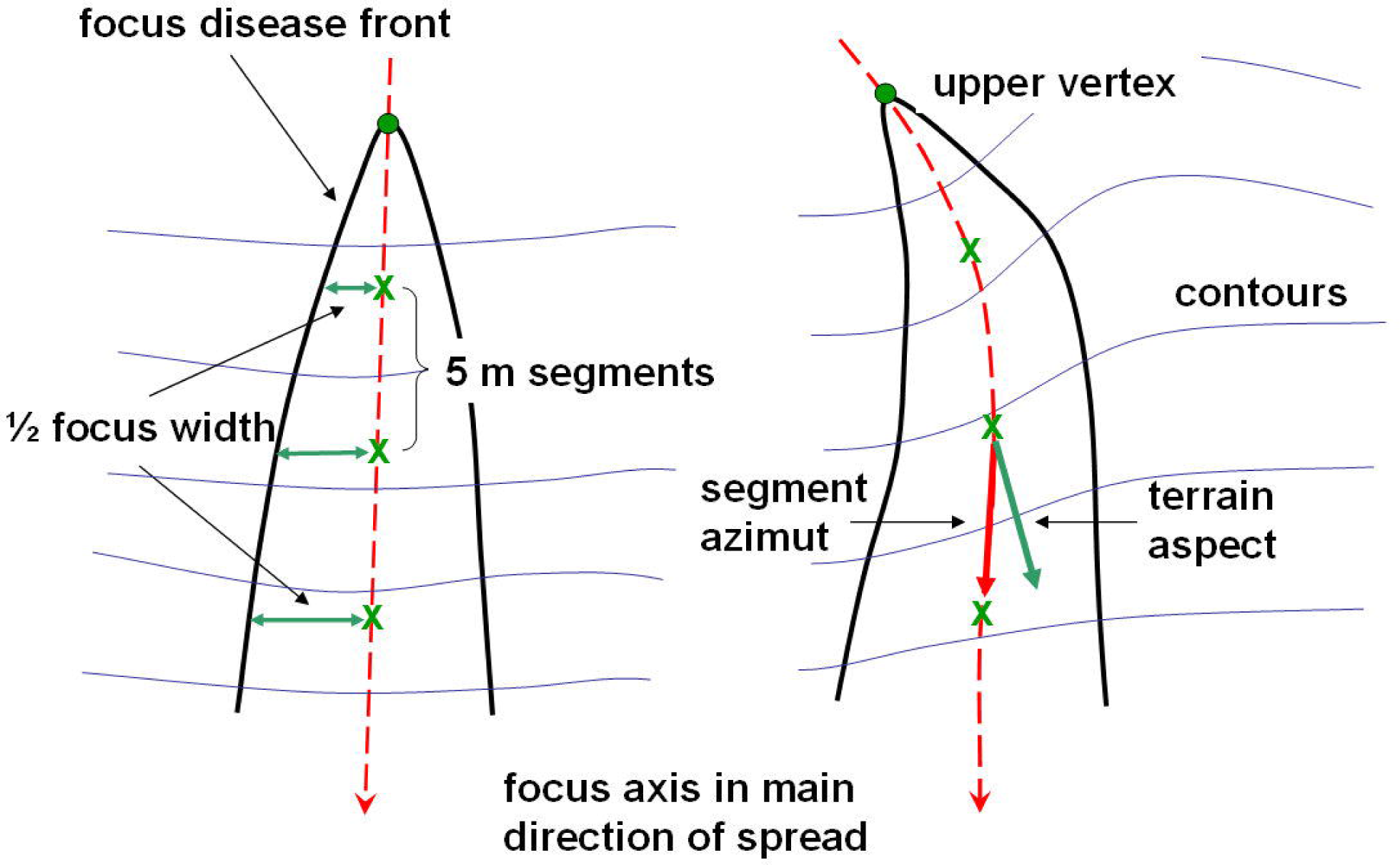
**Descriptive sketch showing measurements taken on foci of disease caused by *Phytophthora cinnamomi* in *Erica umbellata* heathland**. The focus axis along the main direction of spread (red dashed line) was divided in 5m segments (green crosses) to measure focus width and axis azimuth.

Likewise, since the focus width appeared also to increase downslope (Fig. 3), a linear model was used to explore this pattern. In this framework, the horizontal semi-distance between opposite disease fronts (Fig. 4) corresponding to a given focus was considered to be the independent variable. The distance to the upper vertex of the focus was used as the main effect. The terrain slope and curvature were included in the model to test geo-morphology effects on focus width. A linear model was fit by maximum likelihood methods using ‘R’ statistical software version 3.1.1 [56].

### Dispersal pattern of foci

As the aerial imagery revelled, the number of foci increased over time (Fig. 5). Then, the total diseased area and the number of patches present in the field were measured in all images taken over the years to understand how the disease progressed. Several nonlinear models (exponential, monomolecular, logistic, Gompertz, and Weibull) were compared to describe the relationship between accumulated foci number and affected surface over time. The fit of the curves to data was checked both by plotting quantities and by fitting nonlinear statistical models by means of ‘nls’ package on ‘R’ [55, 57].

**Fig. 5.**
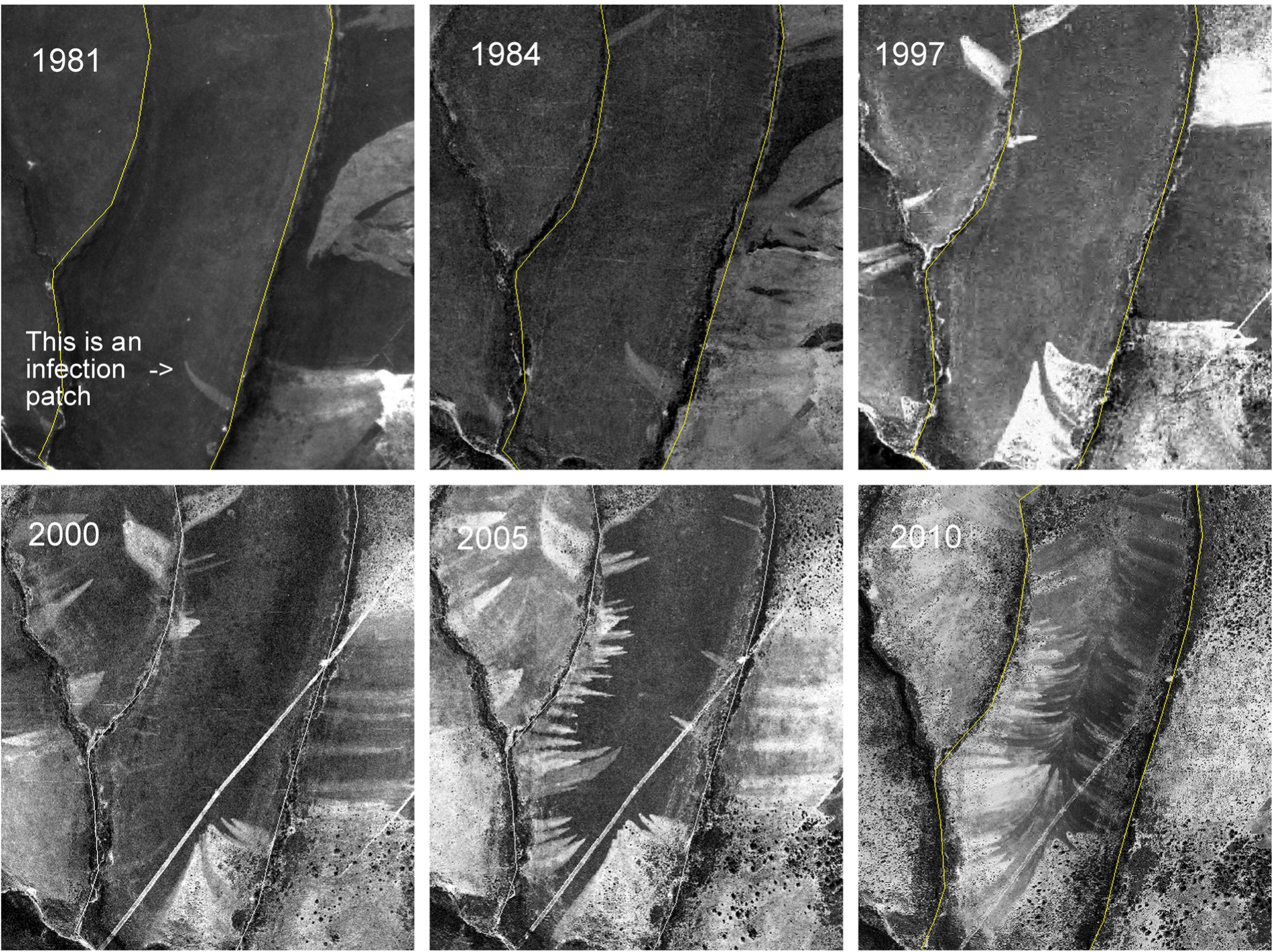
**Aerial images showing proliferation of disease patches on a heathland (*Erica umbellata*) caused by *Phytophthora cinnamomi***. Year of image acquisition is showed in the upper-left corner. The wide patches on the left side of the 1981-1997 images are ploughed plots. Later, in 2000, a firebreak (the diagonal white line) was built. No disease foci emerging from these features were observed. Printed under a CC BY license, with permission from Junta de Extremadura CICTEX, original copyright: Orthoimagery CC-BY 4.0 CICTEX, Junta de Extremadura.

To gain insight into the creation mechanism of new foci, the spatial pattern of the first 79 separated disease patches found in 1984 and 1997 images was analyzed. Based on the previous analysis of the foci growth patterns, the upper vertex of each focus was considered as the nearest identifiable point to the inoculum introduction site (IIS) and used as a reference to describe its evolution. Geographical coordinates of putative IIS were recorded and used as input for ArcGIS in the form of a point data set. To test if IISs were clustered, the observed point pattern was compared to a complete spatial randomness (CSR) Poisson point process of the same intensity. A Nearest Neighbour Index (NNI) defined as the ratio of the observed mean distance to the expected mean distance of a hypothetical CSR process was calculated using ArcGIS [53, 58]. It is considered that the pattern exhibits clustering if the index is less than 1 (the null hypothesis of CSR). The statistical significance of NNI was measured by z-score and p-value.

The cumulative distribution functions (G(r)) of observed and theoretical distances were plotted together and graphically compared to discover at what distances clustering appeared. The Kaplan-Meier estimator of G(r) was used to reduce the bias due to edge effects [59]. To assess the goodness of fit, 39 Monte Carlo simulations (significance level at 0.05) of CSR were carried out and the resulting envelope around the theoretical distribution function was compared with the observed one. To determine where the birth of new foci was more frequent, a smoothed density map (Fig. 6) was created using a quadratic kernel function of ArcGIS described in Silverman (1986, p. 76, equation 4.5, [60]).

**Fig. 6.**
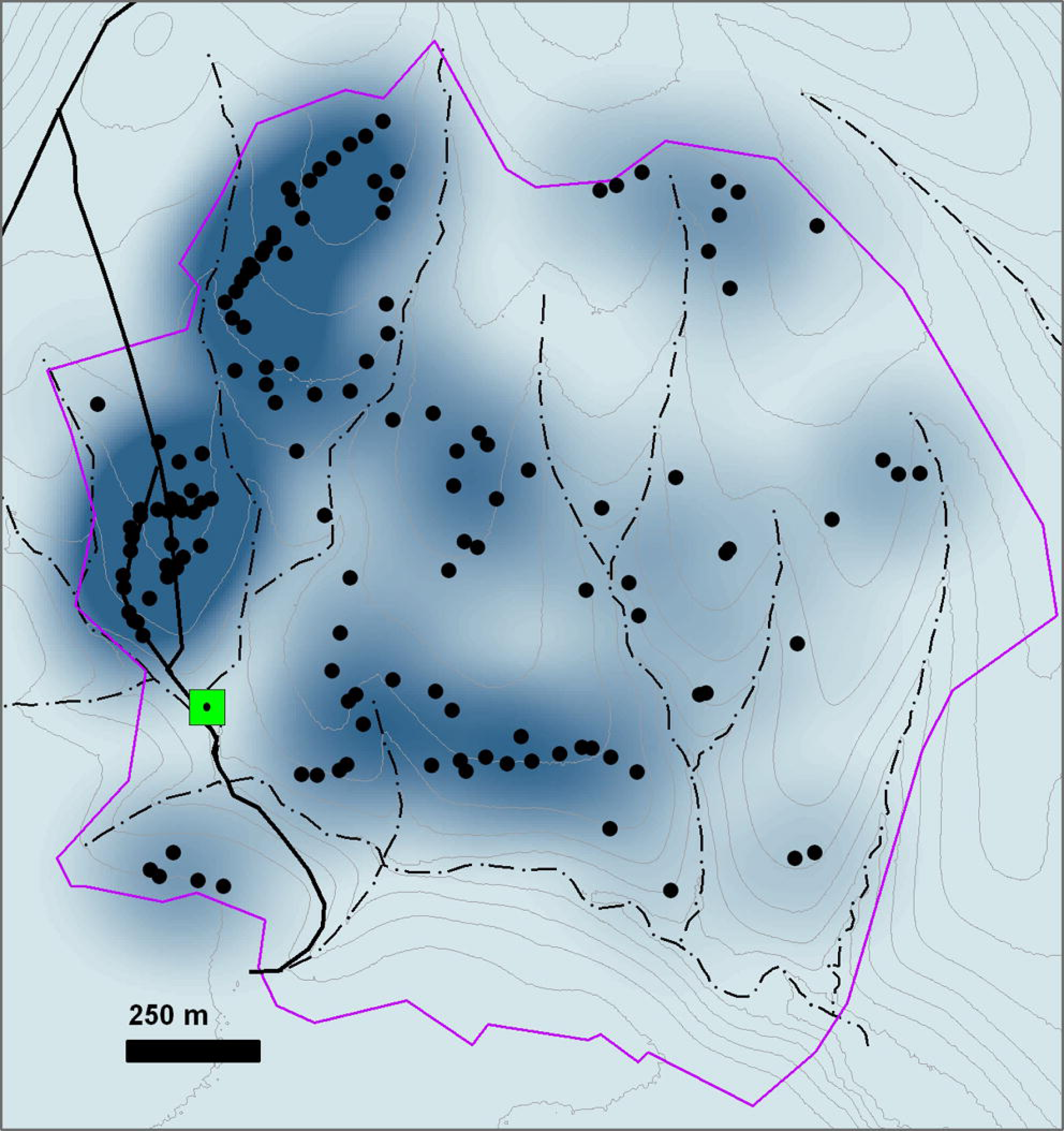
**Map of *Phytophthora cinnamomi* inoculum introduction sites (IIS).** Data from imagery of a heathland stand of Erica umbellata from 1981 to 1997. Legend is as follows. IIS: black dots. Double solid black lines: roads; green square: goat stable and water supply; discontinuous blue lines: streams; brown continuous lines: contour lines; and finally, the shadowed blue background show the IIS kernel point density, the darker the shadow, the higher the density. Purple line shows study area limits. The map has been created in QGIS by the authors using their own data.

To investigate whether topography or terrain morphology are related to the appearance of these new disease foci, a retrospective case-control study based on logistic regression analysis was undertaken. An inoculum source-transport-sink mechanism was hypothesized in order to select risk factor candidates. The diseased area observed in the image taken in 1981 was considered as the inoculum source; the goat flock browsing in the farm was suspected to be the main inoculum transport vector subject to differential habitat preferences; and it was supposed that the soil at each site has different capacity for pathogen establishment.

The set of cases consisted of 79 IIS observed in 1984 and 1997 aerial imagery, while a sample of 173 points generated randomly by ArcGIS inside asymptomatic areas populated by heaths was assumed to be the control set. Risk factor candidates (Table 2) were taken to be variables describing inoculum pressure (foci density and distance to the nearest focus of 1981), topography (distances from streams, stable, rye forage crop, and roads), geomorphology (slope, aspect and plan curvature of the terrain), and proxy variables related to soil water content and temperature (topographic wetness index and spring potential radiation). Each variable was calculated for every 5x5 m cell of a data grid. The cell values of these raster layers at the IIS and the locations of control points were transferred to the data set.

A univariate binomial model with a logit link function was fitted using deviance as a measure of the goodness of fit to identify possible risk factors (‘glm’ package on R software, [56]). The likelihood-ratio Chi-square test was calculated as twice the difference of the log-likelihoods between the model with the risk factor and the null model without the effect. Odds ratio (OR), a measure of association between the exposure factor and the disease outcome, and its 95% confidence interval were estimated. When OR > 1, the factor is associated with higher odds (risk) of disease, if OR < 1, an increase in the factor levels tends to reduce the disease odds, and when OR = 1, the factor does not affect the odds. Univariate effect graphics were performed using the package, ‘popbio’ on ‘R’ software [56, 61].

Finally, to explore the predictive power of these topographic traits, a multivariate logistic model was applied tentatively. Firstly, multicollinearity was examined by computing the correlation matrix and the variance inflation factor (VIF) (‘car’ package, [62]) to avoid redundancy between predictors. Secondly, no redundant variables which, according to univariate analysis, were significantly associated with the onset of new disease foci were used as input for a multivariate analysis aimed at obtaining the most adequate minimal model. Variables were selected using a stepwise process based on Akaike information criterion (AIC) implemented on package ‘MASS’ [63]. To evaluate the predictive power of this model, the receiver-operating characteristic (ROC) curve and the area under the ROC curve (AUC) were calculated (‘pROC’ package, [64]). A model with perfect discrimination has an area of 1.0, as opposed to an assigned value of 0.5 for a model lacking any discrimination power. The Pearson residuals of the model were plotted to identify non-linearity, outliers, leverage and influence. The Spatial autocorrelation of Pearson residuals was tested by means of Moran’s index (‘spdep’ package [65]).

## Results

### Pathogen isolation

The analysis of the Poisson GLM fitted with the data gathered in the field survey shows that the abundance of *P. cinnamomi* depends on position with respect to the disease front (F_(2,15)_ = 13.66, p-value < 0.001). The average number of isolations (2.67 ±0.76, standard error) obtained from samples just outside the symptomatic patch is significantly lower (z = -3.782, p-value < 0.001) than the number of isolations obtained in the front (23.83, ±4.45) or in the bare core of the patch (27.00, ±3.63). No significant difference was found between core and front samples (z = 0.141, p-value = 0.989).

### Diseased foci recognition

Patches of diseased plants were easily detected in all the aerial imagery, although small patches (width < 3 m) were less noticeable on older aerial images. Some of them were identified in field visits or by using the most recent high spatial resolution imagery. As evidenced in field visits, a narrow but sharp disease front between healthy plants and a band of dead heath plants delimited the perimeter of each patch. Field measurements revealed that the average width of this band was 0.46 ± 0.07 meters.

Inside the patches, *Phytophthora cinnamomi* completely destroyed dwarf heath (*E. umbellata*), but scattered individuals of basil-leaved rock rose (*Halimium ocymoides*) and endemic broom *Pterospartum tridentatum* survived. In the inner part of the patches the soil remained bare until a sparse population of gum rockrose (*Cistus ladanifer*) appeared. In some sites, other species such as *Lavandula stoechas* were growing with the gum rockrose. Scattered, 5-10 cm high, dwarf heath seedlings regenerated from the seed bank were observed inside the patches, although they did not survive for a long time. In older patches, pines seedlings (*Pinus pinaster*) appeared in the core, while seedlings of asymptomatic *Arbutus unedo* and *Pyrus bourgaeana* were observed near streams. In one of the oldest foci, initiated more than 30 years ago, poles of maritime pine were present and asymptomatic mature (30 cm high) individuals of dwarf heath were part of the shrub understory community.

### Focus growth patterns

In shape, small patches (< 3 m width) were circular on flat areas, but elliptical on slopes (Fig. 1). More developed patches resembled parabolic shapes oriented downslope (Figs. 1 and 3). These shapes appeared truncated by drainage lines (Figs. 1 and 2), that is to say, no progress of the front disease was observed in any case on the opposite shores of streams. In developed patches, a vertex point of maximum curvature located in the upper and narrow part of each focus was evident (Fig. 3). From the vertex, two front branches forked downslope.

Results of the one-way ANOVA (F_(2,75)_ = 10.91, p-value < 0.001) indicated that the observed rate of focus growth was direction dependent. At early stages, before foci reached streams, the average advance downslope (17.1 ±5.00 m/year) was significantly greater (t = -4.140, p < 0.001) than the spread upslope (0.16 ±0.05 m/year) or laterally, following contours (0.96 ±0.17 m/year).

As confirmed by the regression model (F_(3,138)_ = 158.2, p-value < 0.001, adjusted R^2^ = 0.77) the patch width increases downslope in a parabolic fashion (Table 3). Moreover, terrain curvature and slope had a significant effect on focus width. Wider patches were associated with divergent terrain shapes (spurs) and the narrower with convergent gullies, while increasing slope tended to reduce focus width.

**Table 3.** **Phytophthora cinnamomi focus width analysis.** Linear model to test a parabolic trend on Phytophthora cinnamomi focus width increase with distance from the upper vertex and effects of terrain curvature and slope. Factors coefficients (β), standard errors (SE β) and significance are shown

| Predictor | $\beta$ | $SE \beta$ | t-value | p-value |
| --- | --- | --- | --- | --- |
| Intercept | 0.417 | 0.344 | 1.210 | 0.228 |
| Root square of distance to vertex (m) | 1.091 | 0.050 | 21.617 | <0.001 |
| Plan curvature index | 3.713 | 0.633 | 5.865 | <0.001 |
| Slope (%) | -0.109 | 0.017 | -6.378 | <0.001 |

As for the direction of prevailing growth, the regression fit (F_(1,125)_ = 6600, p-value < 0.001) showed that the main direction of the focus expansion followed the terrain aspect (Table 4). The azimuth of each of the segments into which the main axis of each path was divided could be accurately predicted by the terrain aspect measured at each segment upper point (adjusted R^2^ = 0.98).

**Table 4.** **Phytophthora cinnamomi focus spread direction analysis**. Regression model analysis of the association between focus main axis azimuth and terrain aspect. Factors coefficients (β), standard errors (SE β) and significance are shown

| Predictor | $\beta$ | SE $\beta$ | t-value | p-value |
| --- | --- | --- | --- | --- |
| Intercept | -0.307 | 3.057 | -0.10 | 0.261 |
| Aspect (degrees) | 1.011 | 0.012 | 81.24 | <0.001 |

### Dispersal pattern of foci

Regression analysis showed that the accumulated number of disease foci has been increasing exponentially over time (Fig. 5). Although fits based on logistic and Gompertz models perform relatively well, their characteristic asymptotic growth was not observed in the data. The exponential model (F_(1,4)_ = 167.6, p-value < 0.001) explained 97.1% of the total variation in the accumulated number of foci over time. The estimated apparent infection rate was 0.145 ±0.011 new foci per number of already existing foci and year. In addition, the exponential model (F_(1,4)_ = 231.3, p-value < 0.001) explained 97.9% of the variability in the accumulated diseased area, and a similar growth rate was observed (0.150 ±0.010 ha per ha and year). The total diseased area and the accumulated number of foci turned out to be highly correlated (r = 0.995) due to a low variability in the area per focus (0.102 ±0.003 ha/focus).

The distance methods used to investigate the distribution of inoculum introduction points (IIS) suggest a clustered pattern. The average nearest neighbour distance between IISs (49.0 m) was significantly (NNI = 0.70, Z-Score= -5.2; p-value < 0,001) lower than the expected mean distance (70.2 m), corresponding to the null hypothesis of a completely random spatial distribution of points. Moreover, this clustered pattern occurred in a wide range of reference distances (20-90 m), as could be deduced from the graphical analysis of the observed cumulative distribution and a theoretical CSR cumulative distribution of IIS nearest-neighbour distances. The observed distribution lay well above the typical range of values (Montecarlo envelope) of the G(r) function for a completely random pattern.

Two patterns in the distribution of the IIS could be distinguished visually in the kernel density map (Fig. 6). To the west, two elongated (about 800 m long) hot spots running south to north were around farm access road and tractor tire tracks. The majority of these points were already observed in the 1981 image (Fig. 2). In the rest of the farm, most IISs originated after 1981 and appeared more disperse and random. Nevertheless, evidence for some clustering can be observed.

To analyze the influence of topographic effects on the distribution of foci, fourteen variables were tested individually. Nine of them turned out to be relevant for the probability of a new infection onset during the observation period (Table 5 and Fig. 7). As expected, inoculum pressure, described by the number of foci per unit area that already existed prior to the study (foci density), was a relevant parameter. According to this univariate analysis, sites with low visibility (topographic openness OR = 3.37) or steep slopes (OR = 0.93) were less prone to disease. In contrast, disease occurrence was more probable on sites close to the forage crop or the goat stable. This analysis also shows that heathland was more prone to disease when located on terrains with higher plan curvature (OR = 1.23), in zones elevated from streams (OR = 1.03) or in sites that received more radiation (OR = 1.03).

**Table 5.**
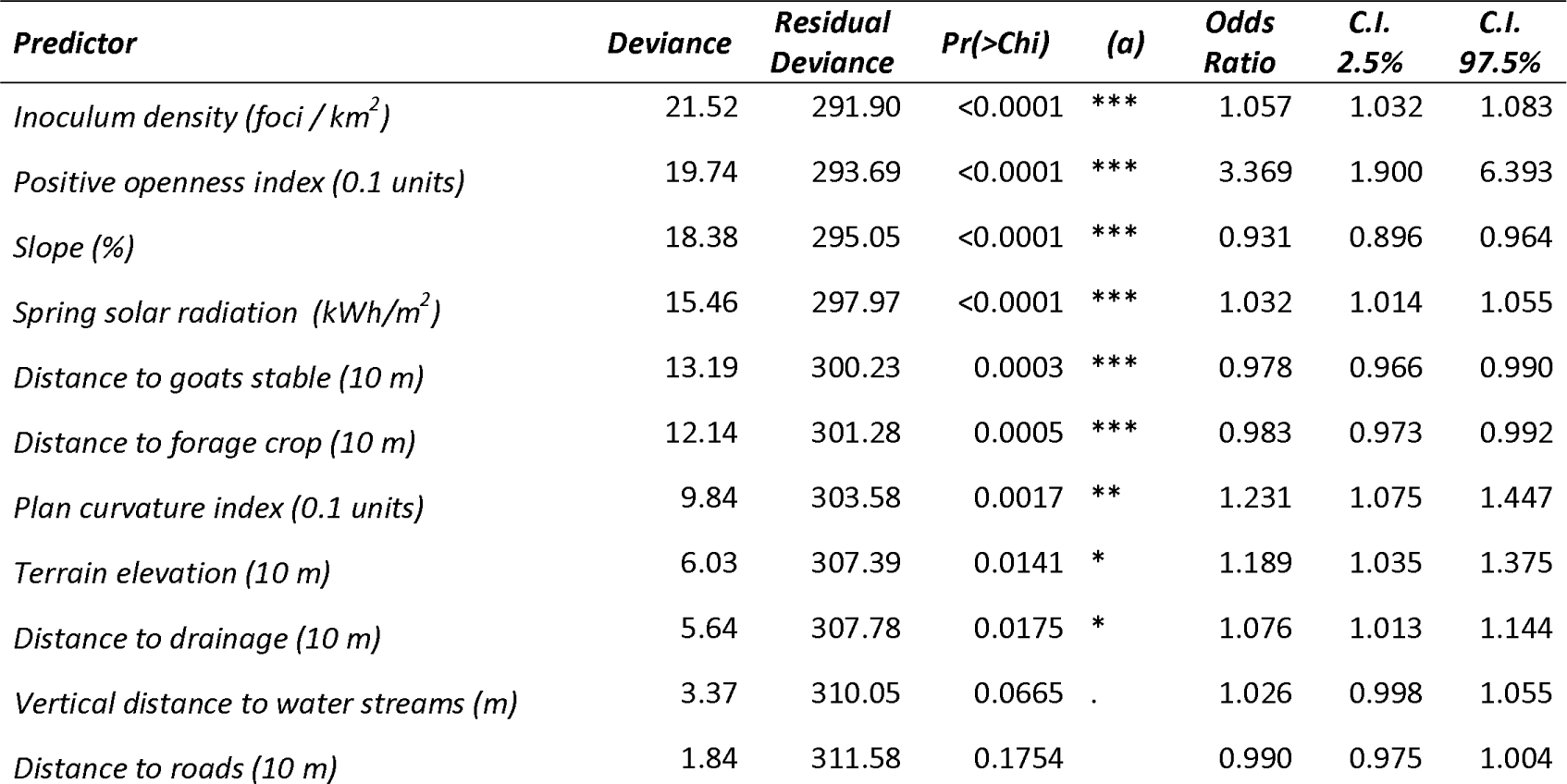
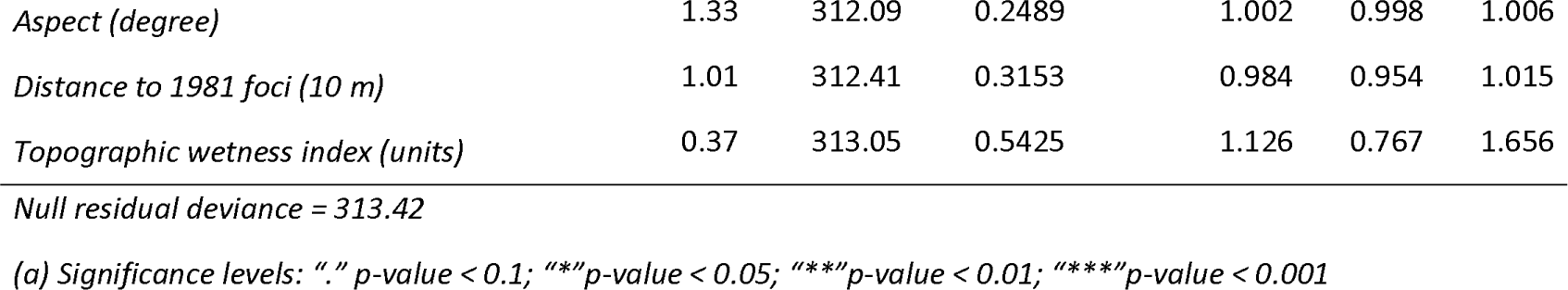
**Univariate risk factor analysis.** Deviance, likelihood ratio test significance and odds ratio for each topographical attribute used as a predictive factor in univariate logistic regression models. As binomial outcome, cases consisted of new infection occurrences and controls were random sites that remained healthy. The units for which the odds ratio was calculated are indicated in parentheses. C.I. = confidence interval (95%).

**Fig. 7.**
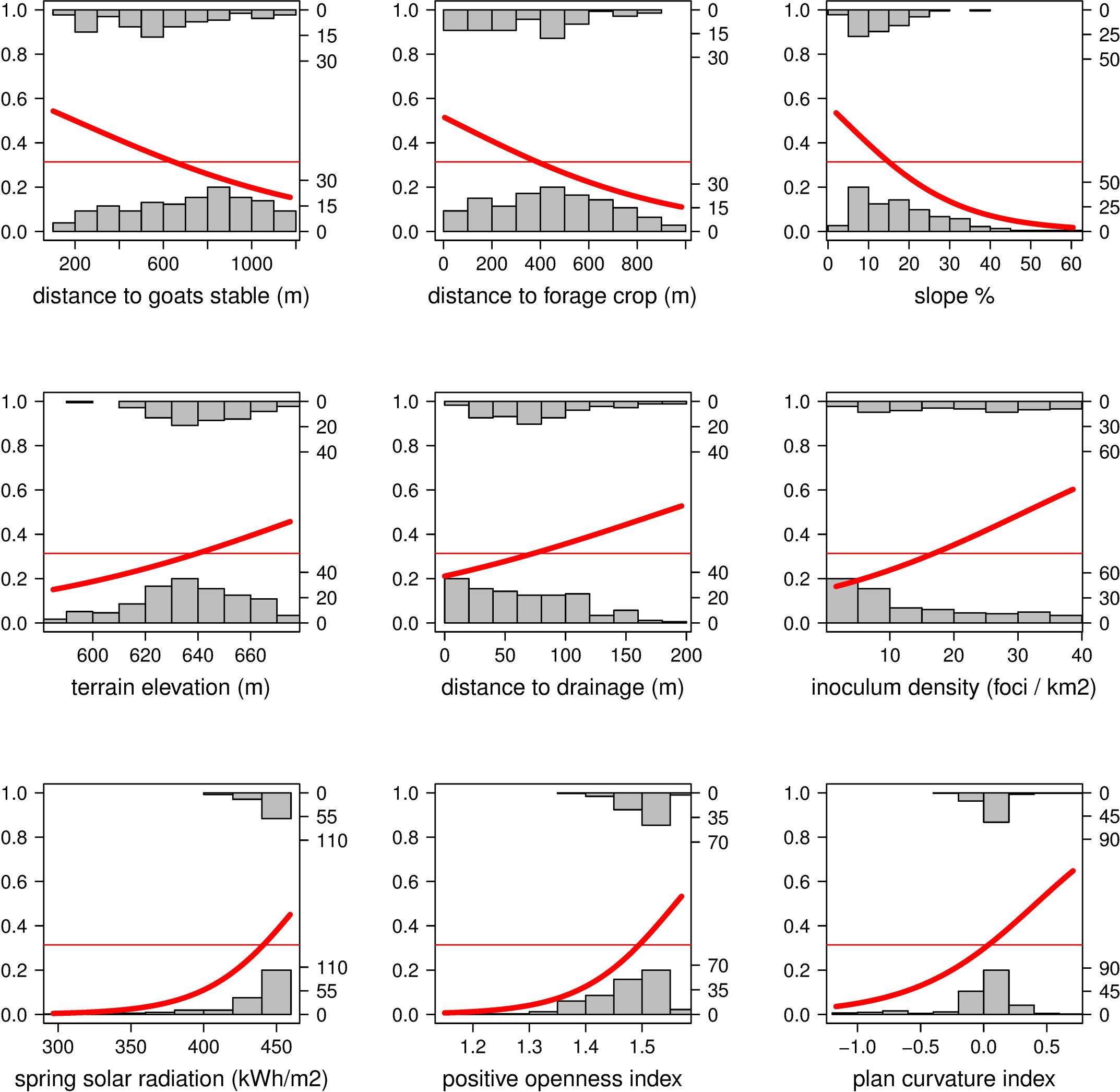
**Univariate effects of risk factors analyzed.** Fitted univariate logistic regression models for topographic factors with a significant effect on *Phytophthora cinnamomi* disease probability. The histograms represent the observed data, in the top are the number of cases (diseased) and in the bottom the number of controls (healthy). Frequencies of both can be read from the on the left Y axis. The red thick line is the predicted probability for the onset of a new *P. cinnamomi focus*. The thin red line corresponds to the average probability without the effect of any risk factor. Both probabilities can be read from the right Y axis. Positive openness and plan curvature are dimensionless numbers without units.

A multivariate analysis was done to fit a minimal adequate predictive model starting with the significant variables of the univariate analysis. No detrimental collinearity between these explanatory variables was detected, with the largest VIF standing at 7.1. A logistic model with four explanatory factors (Table 6) fitted significantly better (model deviance = 44.04; d.f. = 4; Pr(>Chi) < 0.001) than the null model (null deviance = 313.42; d.f. = 251). According to the fitted model, the odds (risk) of a new disease patch onset decrease with increasing distance to stable or forage crop. This decrease was around 2% (OR = 0.98) of the odds at each 10 m distance increment. The model also showed that the more visibility a location offers (animals can see further), the greater the odds (OR = 2.22 for a topographic openness index increase of 0.1 units). Plan curvature also had a significant effect. Spurs or sites with divergent flow across terrain surface were more prone to disease foci emergence than gullies or locations with convergent flows (OR = 1.21 for a curvature index increase of 0.1 units). As a predictive tool, the accuracy of this minimal model was fair, yielding an area under the receiver operating curve of 0.74. No spatial autocorrelation was observed in the model residuals (Morans’I = -0.005, p-value = 0.554). Based on the logistic model a risk map (Fig. 8) for the occurrence of new *Phytophthora cinnamomi* focus was created in ArcGIS.

**Table 6.** Summary of the minimal adequate model for new disease manifestation prediction. After dropping non significant or correlated factors, this logistic model just includes the minimum number of variables witch significantly yield the maximum reduction of the deviance. Estimates and their standard errors are in logits. Odds ratio was calculated for the factors units which are indicated in parentheses. C.I. = confidence interval (95%).

| Predictor | Coef. $\beta$ | SE $\beta$ | Wald z | P>\|z\| | (a) | Odds ratio | (2.5%) | (97.5%) |
| --- | --- | --- | --- | --- | --- | --- | --- | --- |
| Intercept | -10.658 | 5.534 | -1.926 | 0.0541 |  | 0.000 | 0.000 | 0.864 |
| Positive openness index (0.1 units) | 0.796 | 0.363 | 2.192 | 0.0284 | * | 2.217 | 1.115 | 4.652 |
| Plan curvature index (0.1 units) | 0.193 | 0.093 | 2.084 | 0.0372 | * | 1.213 | 1.022 | 1.471 |
| Distance to forage crop (10 m) | -0.020 | 0.007 | -2.780 | 0.0054 | ** | 0.980 | 0.966 | 0.994 |
| Distance to goats stable (10 m) | -0.021 | 0.006 | -3.643 | 0.0003 | *** | 0.979 | 0.968 | 0.990 |

(a) Significance levels: "." p-value < 0.1; "\*" p-value < 0.05; "\*\*" p-value < 0.01; "\*\*\*" p-value < 0.001

**Fig. 8.**
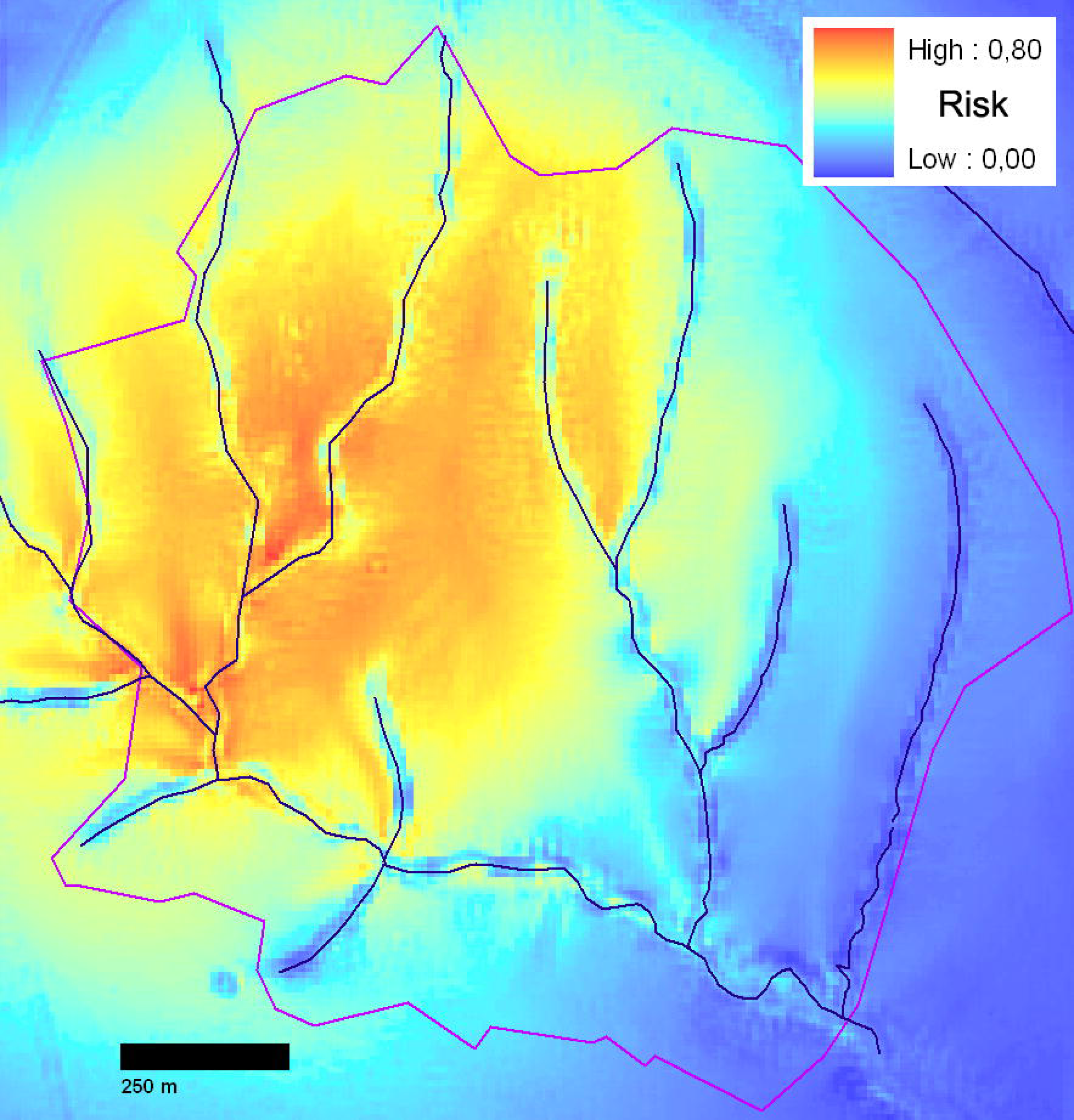
**Risk map for the occurrence of new *Phytophthora cinnamomi* foci in the study area.** The map shows probabilities ranks estimated by a logistic model with four risk factors: topographic openness index (visibility), plan curvature of terrain and distances to forage crop and goat stable. The map has been created in QGIS by the authors using their own data.

## Discussion

In this observational study, we show a clear picture of the spatial dispersion patterns of the disease caused by *P. cinnamomi* at a specific site in Spain. A stand populated by a continuous canopy of a small, highly susceptible heath along with high-resolution imaging provide an invaluable opportunity to observe the spread of the disease and to significantly narrow the location of the initial inocula triggering the formation of disease foci. We focused on the characterization of the shape distribution of seamless patches of dead heaths and their spatial spread over time. Firstly, we found a few small circular or slightly elliptical incipient patches which were mainly located on low gradient slopes. Some developed into largely elliptical or narrow parabolic shapes. These parabolas, which were the most abundant shape observed, extend their branches downslope with their trajectory curving to follow a fairly right angle to the contours. Growth was more pronounced in their lower parts, where the parabola branches open up in a fan-like pattern. The longitudinal growth of such patches was interrupted whenever a stream was encountered. The statistical analysis confirmed that parabola direction and amplitude were largely determined by topographical features, namely terrain aspect and plan curvature.

Secondly, we found the number of heath mortality patches increased exponentially over time and a correlated spatial pattern emerged. In this case, the spatiotemporal evolution was less clear-cut than at the level of single foci, although foci clustering could be proved. The case-control study revealed that nine out of the fourteen topographic factors examined were associated with the probability of observing disease emergence in a given site. Although the emergence of foci was mainly related to the spatial density of former foci, that is, with abundance of inoculum sources, other predictive variables were also found to play a role. The probability for the birth of a new focus was found to depend significantly on distances to flock resting or feeding sites, terrain visibility, slope, and soil temperature or humidity proxies such as radiation income or distance to water streams.

The circular patterns of the incipient patches observed on flat areas could be attributed to the effect of a slow rate radial growth based on a root-to-root contact, mycelium growth, and limited zoospore movement, as described by Hill et al. [26], Wilson et al. [27], and Shearer et al. [28] in Banksia woodlands. This propagation in concentric circles could be described by the Skellam diffusion-reaction model in its more simple form [33].

However, in the case of the parabolic shapes observed in the Villuercas slopes, other dispersion forces appear to be at play. It is very plausible that the action of the water flowing through the soil, as observed by Shea et al. [3] and later by Kinal et al. [29], may play an important role as a second vector here. Thus, the combined action of both dispersion vectors could possibly be enough to explain the parabolic shapes of these foci. From a more mathematical point of view, the above hypothesis appears to fit reasonably well into a picture based on the advection-diffusion-reaction model proposed by Shigesada and Kawasaki [44].

As far as the lateral growth of the foci is concerned, our findings can be summarized as follows. Firstly, in the circular patterns corresponding to early stages of the growth process, we observe geometry-preserving radial growth. Secondly, we find that the rate of spread of well-developed, parabolic foci depends on the distance to the vertex (Table 3). Finally, we find a remarkable effect of the topographic curvature and slope on focus growth. That is, the terrain morphology could be contributing to concentrating or dispersing propagules by altering water flows and thereby affecting the amplitude of the parabolas. A consequence of this is that the final area occupied by a disease focus can be accurately estimated from the position of the inoculum introduction point, the time since settling, the geomorphology and the distance to water drainage (which acts as a natural barrier for the disease front).

Despite the above, the most important contribution to the total affected area in the stand was the formation of new infection foci, whose number was found to grow exponentially. Based on the morphology of the disease foci, it was possible to estimate the location of the inoculum introduction point of each focus with confidence. Then, we hypothesized that transport of propagules from diseased areas to distant healthy sites in the stand could be mainly attributed to the goat flock, and that the behaviour of these animals is partly conditioned by topographic traits. We also hypothesized that the probability of establishment of the *Phytophthora* inoculum depends on soil suitability, which again may depend on topographical proxies. This more complex situation of dual or stratified dispersal via water in the soil and the motion of animals could possibly be addressed by means of a diffusion-reaction model with two different channels of dispersal [43].

Although in the scope of a case-control study it is difficult to infer well-established causal relationships, it is possible to put forward some factors that are likely to play a role in short- and long-distance dispersal. For example, one expects a large number of propagule depositions in sites where disease density is higher or in sites which are more frequently visited by animals. In contrast, a smaller number of discharges of infected soil particles would be expected in areas which animals are reluctant to visit. As pointed by Bailey et al. [39] and Lachica et al. [38] in the case of goats, locomotion over the steepest slopes requires an extra energy expenditure that animals tend to avoid. Similarly, less preferred sites also tend to be habitats where animals have a short range of vision, such as in areas with low topographic openness, e.g., closed gullies. Indeed, reduced visibility results in a decrease of foraging efficiency because of the longer time needed to detect threats and because of the natural tendency of animals to feed close together increasing foraging competition [40].

As far as site suitability to pathogen establishing is concerned, the univariate logistic models show that the probability that *P. cinnamomi* gives rise to a new infection patch grows significantly with increasing amounts of sun radiation received at a site during spring, with increasing distance from streams, and with increasing positive curvature. All of these indicate that drier sites were more prone to successful inoculum settling. This is in agreement with the findings of Wilson et al. [27], who reported that a sun index of a similar order of magnitude favoured the presence of *P. cinnamomi* in the sclerophyll vegetation of south-eastern Australia. On the other hand, it is in apparent disagreement with studies by Vannini et al. [9] and Balci et al. [30] which point to a greater pathogen presence and disease incidence in humid valley bottoms near streams.

However, our conclusions so far refer to the location of the inoculum introduction point, not to the total affected area. Focusing on the latter, we also found that a greater area was affected in valley bottoms near the streams of the Villuercas site due to the fact that disease foci increased their width downslope. A possible explanation to the fact that IIS appeared farther from streams is that *Phytophthora* transport needs sticky soils gathered by animals on water-saturated zones, but its release and establishment could be more probable on drier, airier and warmer sites. In these locations, mud particles can be released more easily and soils have better conditions for growth of pathogens and plant roots during the rainy season. The proliferation of *P. cinnamomi* is limited by both temperature and water availability, so if spring conditions of the soil are relatively wet across all the site, temperature may well be the most important restrictive factor [13].The fact that the emergence and growth of disease foci caused by *Phytophthora cinnamomi* could be determined to a certain degree by topographical traits has relevant implications for the management of the disease. Apart from a better understanding of disease spread mechanisms, one important consequence is that the disease front location could be modelled and used in more opaque ecosystems to improve detection sampling schemes, to trace more accurate containment buffers around diseased vegetation and to select target areas or plants in treatment programmes. Even so, difficulties to locate inoculum introduction points will arise where inapparent or latent infections takes place, in these cases putative initiation sites could be marked upslope of symptomatic vegetation.

In our specific case study, it was shown that the major contribution to the increase in the diseased area stems mainly from new foci creation, whereas the enlargement of already existing foci plays a less relevant role. This is in agreement with the accelerated spread of the disease described by a stratified dispersion model [44]. Understanding and predicting how the pathogen is carried from one site to another is critical to abate the epidemic. In this context, the present study shows that topography may have an important influence on the underlying transport processes.

The present study points to three elements upon which dynamic models, such as those proposed by Thompson et al. [14], Shigesada et al., [44] or Okubo and Levin [33], can be developed for *P. cinnamomi* spread: 1) the growth of existing foci is controlled by a combination of diffusive root-to-root contact and advection by water flows, 2) the formation of new foci is mediated by long-distance transport due to the motion of animals and humans along certain preferential pathways, and 3) in both cases, topography should play a major role, that is, certain sites are more frequently visited by animals than others, and water always flows downhill. In view of the above, incorporating the effect of topographical features in the framework of sophisticated dynamic models may pave the way for further advances in the theory of reaction-diffusion processes underlying *P. cinnamomi* propagation.

Finally, our study also shows that models taking advantage of low cost and highly accurate terrain elevation data can be used to create risk maps and to test new environmental scenarios or management tactics. Nevertheless, it should be acknowledged that the patterns described here were observed in a monospecific ecosystem populated by a continuous cover of small and highly susceptible species living in a particular habitat, soil and topography. Therefore, extrapolation to a more complex or different ecosystem with a higher diversity of species, susceptibilities, vegetation coverture, soil conduciveness or different animal vectors is not straightforward and further research is needed.

## Acknowledgements

The authors thank Ramón Santiago and the forest ranger Juan Carlos Herrera for their support during the field surveys. We are indebted to Pedro Antolín and the Lab Technicians Emi Galván and Eusebio Dorado for their assistance with the pathogen. We also acknowledge the scientific advice of Angel Felicísimo, Mariola Sánchez and the writing assistance of Elena Montero.

## Supporting information

**S1 File. Dataset_brezos**. Excel file containing one spreadsheet with the variables codification and six spreadsheets with the data used in this study

